# ZIPcnv: accurate and efficient inference of copy number variations from shallow whole-genome sequencing

**DOI:** 10.1101/2025.06.13.659496

**Authors:** Zhengfa Xue, Jingyu Zeng, Jiajing Yuan, Tianci Wang, Xin Lai, Lin Wang, Yu Wang, Huanhuan Zhu, Xin Jin, Jiayin Wang

**Affiliations:** School of Computer Science and Technology, Faculty of Electronics and Information Engineering, Xi’an Jiaotong University, Xi’an, Shaanxi, China 710049; Shaanxi Engineering Research Center of Medical and Health Big Data, Xi’an Jiaotong University, Xi’an 710049, China; State Key Laboratory of Genome and Multi-omics Technologies, BGI Research, Shenzhen 518083, China; The Innovation Centre of Ministry of Education for Development and Diseases, School of Medicine, South China University of Technology, Guangzhou 510006, China; Shenzhen Key Laboratory of Transomics Biotechnologies, BGI Research, Shenzhen 518083, China

**Keywords:** Genomics, Copy number variations, Shallow whole-genome sequencing, Zero-inflated, Low coverage

## Abstract

Shallow whole-genome sequencing (sWGS), a rapid and cost-effective sequencing technology, has gradually been widely adopted for CNV analyses. However, with genome-wide coverage of only 0.1-5x, sWGS data exhibit a pronounced zero-inflation phenomenon, whereby a large fraction of loci contains zero sequencing reads. Zero-inflation causes read counts to fluctuate by several-fold between adjacent windows. As a result, random upward blips in coverage can be misinterpreted as copy-number gains (false positives), and true deletions often become indistinguishable from pervasive zero-coverage noise. In addition, existing CNV detection tools developed for sWGS data often struggle to adapt across different CNV sizes. These combined effects severely constrain the accuracy of CNV inference. To address above challenges, we propose ZIPcnv, a novel CNV detection tool specifically designed for sWGS data. First, we apply a large sliding window to smooth the raw read depth signal, which transforms the original zero-inflated statistical characteristics into approximately normal distribution characteristics. We then design a statistical process model that robustly detects persistent shifts under high background noise using a cumulative sum strategy, classifying genomic regions into candidate and non-candidate CNV regions. Finally, dynamic sliding windows are used for one-pass detection of CNVs of varying lengths, with window size adapting to the CNV region size. We evaluated the performance of ZIPcnv on simulated data and 190 real whole-genome sequencing samples. Experimental results show that ZIPcnv consistently outperforms currently popular CNV detection tools. The ZIPcnv source code is freely available at https://github.com/Nevermore233/ZIPcnv.

## Introduction

In some clinical scenarios, besides economic consideration, only trace amounts of cfDNA or ctDNA can be detected in peripheral blood or other body fluids of patients, making high-depth sequencing technically impractical. Thus, shallow whole-genome sequencing (sWGS), which offers a rapid, cost-effective solution for analyzing these low-input samples, has rapidly evolved into a routine technique in genetic diagnoses, as well as many clinical practices **[1–2]**. Current sWGS applications mainly focus on CNV biomarkers, include non-invasive prenatal testing (NIPT) **[3]**, tumor monitoring **[4]** and clonal analysis, liquid biopsy analysis on blood and other effusion fluids **[5-6]**, profiling fresh or frozen tumor tissues **[7]**.

As many CNV detection tools have been developed during the last two decades **[8-10]**, does coverage matter? Unfortunately, yes. Detecting CNVs in sWGS data faces two main challenges: (I) the read-depth distribution becomes warping along with decreasing of sequencing coverage, while (II) poor adaptability to varying CNV scales due to the use of fixed-size window strategies. First, during lowing the sequencing coverage, sWGS data gradually exhibits significant zero-inflation, meaning a large proportion of genomic regions have zero read-depth **[11]**. Such zero read-depth regions are the mixture of copy-number loss regions (true zero regions) and genomic regions which are not sampled in sequencing (fake zero regions). On one hand, real deletion signals are often obscured by background noise from zero-coverage regions, resulting in false negatives. On the other hand, dramatic fluctuations in read depth between neighboring windows affect the statistical means and standard deviation. As a result, random increases in read depth under biased statistical tests can easily be mistaken for copy number gains, leading to false positives. As statistical features are often continuous, most current CNV detection tools assume that sequencing read depths follow classical distributions such as Poisson or Negative Binomial (NB), which are inherently suited to highcoverage data **[12]**. These tools generally achieve good performance at moderately low sequencing depths (above 3X). However, in typical ultra-low coverage scenarios such as NIPT, where coverage is often below 1X, CNV detection tools based on these distributional assumptions perform poorly. Although some CNV tools designed for sWGS data incorporate mixture Poisson models or variance adjustments to account for the dispersion in low-coverage data, they do not explicitly model the abnormally high frequency of zero values **[8,9]**. Consequently, under extremely low-coverage conditions, these tools struggle to distinguish true CNV signals from extensive background noise caused by structural zero-depth regions.

From statistical theory perspective, neglecting zero-inflation fundamentally constitutes model mis-specification, as the chosen distributions significantly underestimate the probability of zero occurrences. Such an oversight leads to a systematic underestimation of the true variance in the data, rendering models overly sensitive to short-term fluctuations **[13]**. As a result, random noise can easily be misinterpreted as genuine CNV signals, while true CNV events may remain obscured by the zero-coverage noise background (leading to increased false negatives). Therefore, zero-inflation is a critical obstacle that undermines the statistical robustness and limits the overall detection capability of CNV analysis methods.

Second, RD-based methods typically require the use of a fixed window size to segment the genome and compute the average read depth within each window **[14,15]**. Ideally, window boundaries should precisely align with the true CNV breakpoints, however, in practice, once window sizes are fixed, it becomes difficult to effectively detect CNVs of varying lengths simultaneously **[16]**. The selection of an appropriate window size is heavily influenced by the research objectives or the scale of CNVs of interest. Larger windows can smooth out the random fluctuations caused by zero inflation in sWGS data, but they also dilute localized, small-scale CNV signals, resulting in false negatives and the loss of clinically relevant variation. Conversely, smaller windows may increase sensitivity to fine-scale CNVs but are more vulnerable to stochastic noise, leading to a higher false positive rate and reduced specificity. Although iterative optimization of window size can theoretically improve detection performance, such procedures are computationally expensive and time-consuming. Scale diversity is a common characteristic of CNV alterations observed in real biological and clinical samples, and fixed-window strategies are inherently incapable of accurately capturing CNV events across different size ranges. Therefore, developing detection tools that can adaptively handle CNV events of varying sizes is crucial for improving the accuracy of CNV detection in sWGS data.

In this study, we propose a novel CNV detection tool for sWGS data, named ZIPcnv. To address the prevalent issue of zero inflation in sWGS data, we apply a sufficiently large sliding window to smooth the raw read depth signal, followed by a log2 ratio transformation. Such a procedure converts the zero-inflated statistical properties of the original data into a statistic that approximates a Gaussian (normal) distribution. Building on this transformed distribution, we employ the cumulative and alarm-triggering characteristics of the cumulative sum (CUSUM) control chart to adaptively adjust the window size according to the scale of candidate CNV regions, enabling accurate detection of true CNV signals across multiple size scales. Experimental results demonstrate that ZIPcnv outperforms existing classical CNV detection tools in terms of sensitivity, precision, and F1 score.

## Materials and methods

The input of ZIPcnv consists of a sequencing alignment file (bam format) and a set of reference baselines (npz format) (Figure 1A). The core workflow of ZIPcnv is divided into three primary steps (Figure 1B and C): (1) correcting base-level coverage, (2) selecting candidate CNV regions, and (3) CNV calling. The output of ZIPcnv is a CNV calling report file (Figure 1D). As a RD-based method, ZIPcnv employs a cumulative sum (CUSUM) control chart **[17]** to identify candidate copy number aberrations and utilizes a dynamic sliding-window strategy to accommodate the detection of CNVs of varying sizes. Prior to running the pipeline, data normalization (Supplement 1) and reference baseline construction (Supplement 2) are conducted.

**Figure 1.**
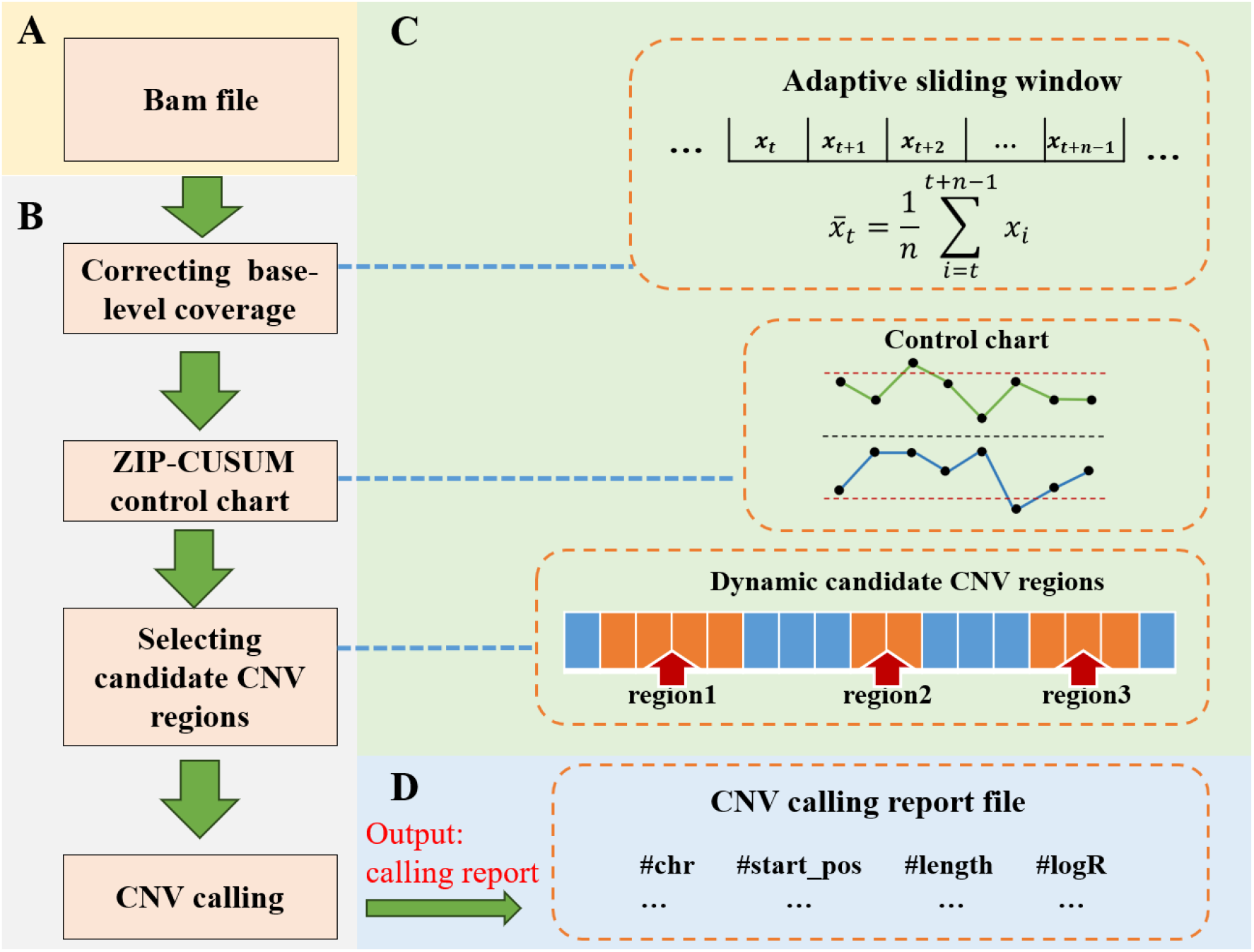
Workflow of ZIPcnv. A. The input for ZIPcnv is a BAM-format sequencing file. B. The core workflow of ZIPcnv consists of three main steps: (1) correcting base-level coverage, (2) selecting candidate CNV regions, and (3) performing CNV calling. C. Specific details of the core steps in ZIPcnv. D. The output of ZIPcnv is a CNV calling report.

### Depth distribution of sWGS data and correcting base-level coverage

In sWGS data, a large number of genomic regions lack sequencing read sampling, resulting in a substantial number of zero read depth values and exhibiting a pronounced zero-inflation characteristic. The initial read depth at single genomic loci can thus be modeled as a mixture process of a zero-inflated Poisson (ZIP) distribution **[18]**—namely, a large proportion of structural zeros combined with a small fraction of non-zero read depths that follow a Poisson distribution. The ZIP distribution is defined as follows:

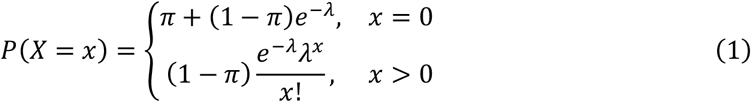

Here, *π* represents the proportion of structural zeros, and *λ* is the rate parameter of the Poisson process. Additionally, the zero-inflated negative binomial (ZINB) distribution **[19]** can also be used to model the locus-level depth distribution in sWGS data. However, the parameters of the ZINB distribution tend to be unstable, and the computational cost is significantly higher.

To achieve more robust modeling under low-coverage conditions, it is necessary to transform the original zero-inflated read depth distribution into a form more suitable for subsequent statistical detection. In this study, we introduce a segment sliding average strategy to smooth the genomic read depth. In this study, we first calculate the base-level depth *x*_*i*_ at each genomic position. Then, a fixed large sliding window is applied to partition the genome sequence into multiple segments. The average number of reads within each segment, denoted as 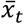, is assigned as the read depth value at the starting position of that segment. The corresponding formula is given in Equation (2):

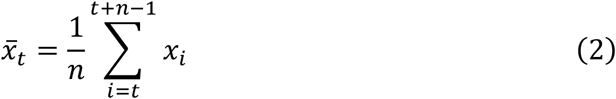

Here, 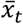 represents the moving average at position *t, n* is the window size (*n* ≥ 3000), and *x*_*i*_ is the base-level depth at position *i*.

According to Central Limit Theorem (CLT), the mean of a large number of approximately independent and identically distributed random variables tends toward a normal distribution as the size of segment increases **[20]**. When the size of the segment is large, regardless of the original distribution of statistic *X*_*i*_, as long as the expectation and variance are finite, the following holds:

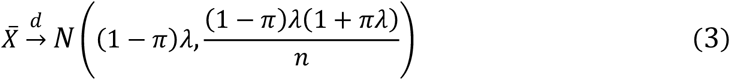

Here, *π* represents the proportion of structural zeros, and *λ* is the rate parameter of the Poisson process, *n* is the size of the segment. An approximately normal distribution not only aligns with the assumptions commonly made in classical CNV detection methods, which assume that sequencing data represent a relatively continuous, symmetric, and stable process, but also provides a solid theoretical foundation for downstream CUSUM-based statistical procedures that rely on the assumption of normality. We have provided the specific details and theoretical proof for the depth distribution transformation of sWGS data in Supplement 3.

### Selecting candidate CNV regions via a dynamic statistical process

#### (I) Construct the cumulative sum statistic

To simultaneously detect CNVs of varying sizes, we designed a dynamic sliding window strategy. Using a cumulative sum (CUSUM) control chart, we identified regions with copy number abnormalities and partitioned the genome into candidate and non-candidate CNV regions. According to CUSUM theory, when (1) the data distribution is approximately normal and (2) the abnormal signal manifests as a weak but continuous shift in the mean, the CUSUM chart exhibits asymptotic optimality in the statistical sense **[21]**. A detailed introduction to the CUSUM model and its theoretical basis for application to sWGS data can be found in Supplement 4. Previous studies have shown that abnormal *log*_2_ copy ratio (*logR*) values across the genome often indicate the presence of CNVs **[22]**. Therefore, we use the *LogR* between observed and expected depth within each large window as the monitoring statistic (or target value) for the CUSUM control chart. Suppose there are *m* monitoring points, and let 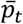 denote the expected depth at the *t*-th site (*t* ∈ {1, 2, 3, …, *m*}) within the control baseline. The target value 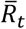 can then be represented as:

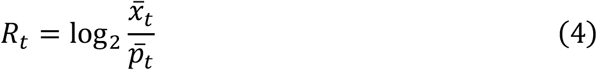

where 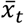 represents the average number of reads within the window. We define *C*^+^ as the cumulative sum statistic representing positive shifts (upward drifts) of the target value *R*_*t*_ (CNV gain), and *C*^−^ as the cumulative sum statistic representing negative shifts (downward drifts) of the target value *R*_*t*_ (CNV loss). The statistics at time point *t* can be expressed respectively as follows:

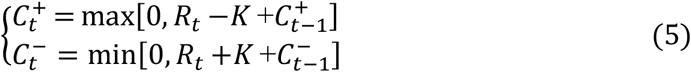

The statistics 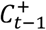 and 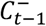 represent the positive and negative cumulative sums at the previous position, respectively, with initial values set to zero.

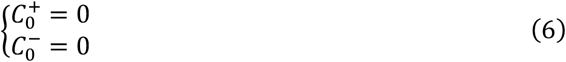

The parameter *K* is the reference value, which controls the sensitivity of detection. Specifically, cumulative sums of the target value *R*_*t*_ begin accumulating only when deviations from *K* occur. A smaller *K* value results in higher sensitivity. In ZIPcnv, the value of *K* is set to 0.3, which aligns with the alert threshold commonly adopted by most CNV detection tools, implying that cumulative sums begin accumulating only when the target value *R*_*t*_ exceeds ±0.3.

We illustrate the workflow of equations **(5)-(6)** through Table 1, which is provided as an illustrative example rather than reflecting actual data. Due to the presence of the maximum and minimum functions, positive and negative deviations (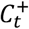 and 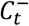) are calculated separately and accumulated independently. Specifically, the maximum function ensures only positive deviations are accumulated, whereas the minimum function ensures only negative deviations are accumulated. Consequently, positive deviations (CNV gain) and negative deviations (CNV loss) do not interfere with each other, enabling the model to clearly differentiate and independently handle duplication and deletion events. When *t* = [1, 2, 3], the target values *R*_*t*_ are within the normal range, and both positive and negative cumulative sums remain at zero. At *t* = [4, 5], the target values *R*_*t*_ exhibit positive deviations, accumulating positive deviations of 0.4 and 0.2, respectively, resulting in a total cumulative deviation of 0.6 at *t* = 5. Subsequently, at *t* = [6, 7, 8], the target values *R*_*t*_ revert to the normal range, and thus the cumulative deviations are reset to zero, indicating the presence of possible noise in that region. As can be seen, ZIPcnv effectively disregards short-term deviations caused by noise or random fluctuations when handling actual copy number variations, thus focusing specifically on long-term trends. Between *t* = 9 and *t* = 13, The 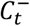 exhibits a clear cumulative downward trend. Even though at *t* = 11 the target value *R*_*t*_ is within the normal range, negative deviations continue accumulating, indicating a potential region of CNV loss.

**Table 1.**
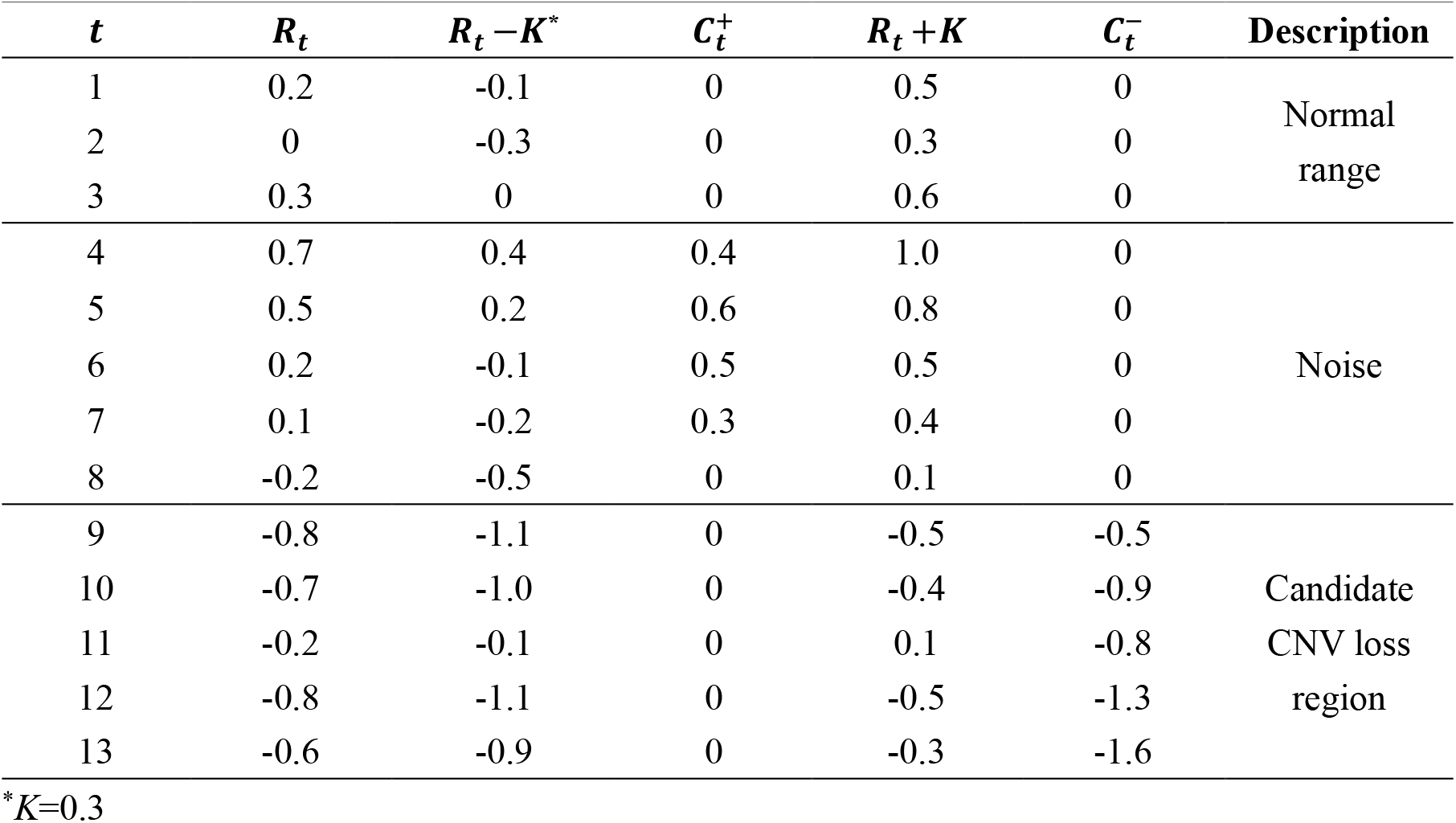
Workflow of equations (5) and (6).

#### (II) Selecting candidate CNV regions

As shown in Figure 2, we use control limits to divide the genome sequence into candidate and noncandidate CNV regions **[23]**. Regions within the control limits are considered non-candidate CNV regions, while those outside the limits are designated as candidate CNV regions. The center line (*CL*) is set to 0, and the upper and lower control limits (*UCL* and *LCL*) are defined as:

**Figure 2.**
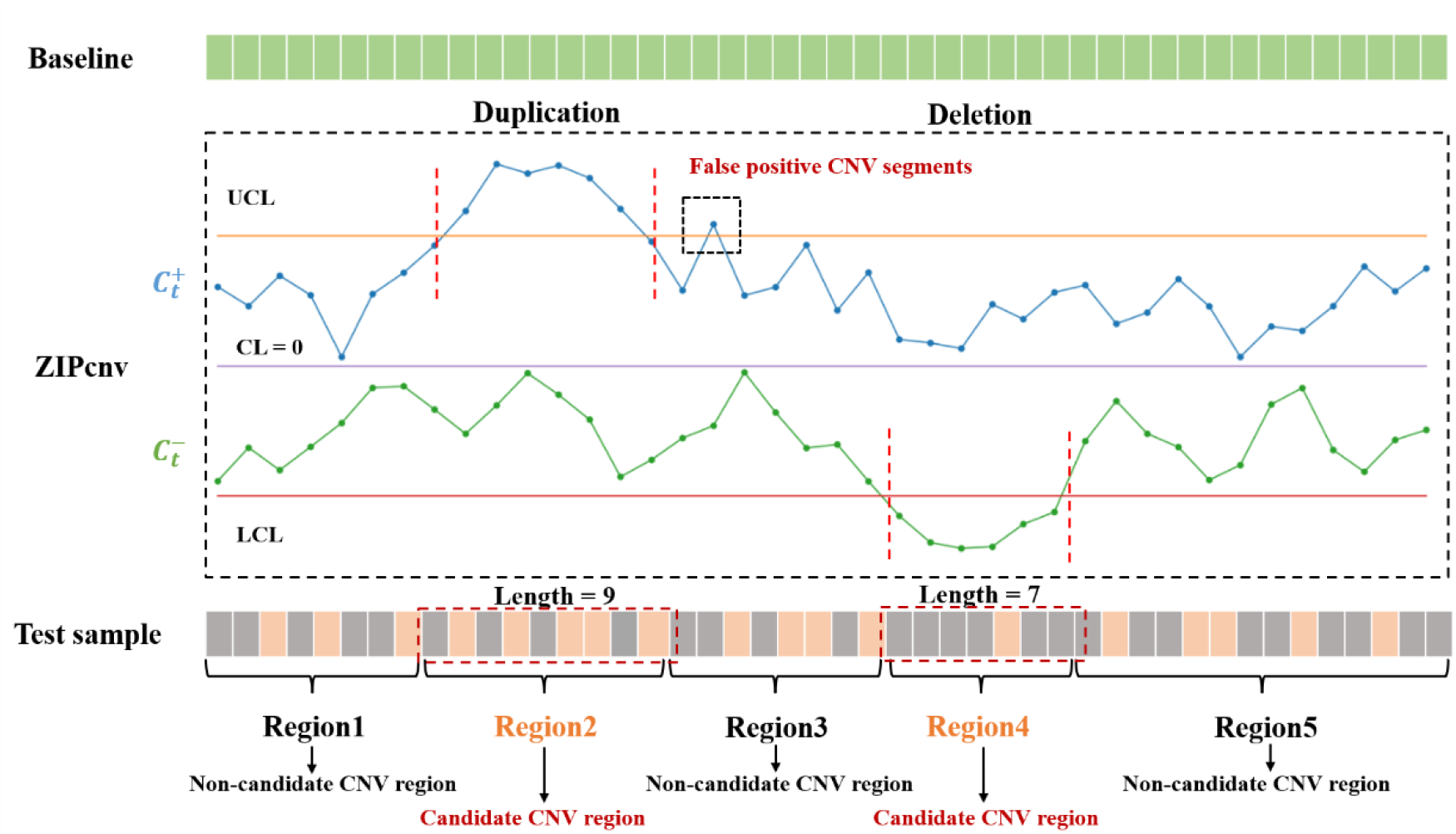
Workflow of the CUSUM control chart in ZIPcnv. When traversing regions 2 and 4, the cumulative sum exceeds the control threshold, triggering an alarm. Window size is then determined based on the alarm region. Similarly, due to noise and zero-depth intervals, the positive and negative cumulative sums may fluctuate and temporarily exceed the CUSUM threshold, generating transient alarms (highlighted by black dashed boxes), which may lead to false-positive events. These intervals are too short to constitute true CNVs, so we filter out any alarm span shorter than 1,000 bp in accordance with CNV definitions.

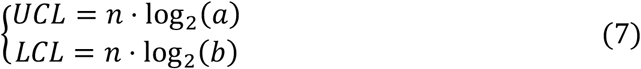

where *n* is the sliding-window size of Equation (2). The factors *a* and *b* in the formula control the UCL and LC L thresholds, respectively. The default values of *a* = 3/2 and *b* = 1/2 correspond to the expected read-depth changes for single-copy loss and gain in an ideal diploid genome. In practical applications, a and b can be adjusted flexibly based on the research objectives: larger values of a or smaller values of b will increase the specificity of the detection, while smaller values of a or larger values of b will enhance the sensitivity of the detection. A positive cumulative deviation 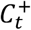 exceeding the *UCL* signals a potential copy-number gain and triggers an alarm; conversely, a negative cumulative deviation 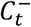 falling below the LCL indicates a potential copynumber loss. As shown in Figure 2, the statistics for regions 2 and 4 exceed the control limits, and we therefore classify these regions as candidate CNV regions. In contrast, the statistics for regions 1, 3, and 5 fall within the control limits and are considered non-candidate CNV regions. Additionally, due to noise, the cumulative sums may briefly exceed the control limits, resulting in transient alerts (highlighted by black dashed boxes in Figure 2). These intervals are too short to constitute true CNVs, so we filter out any alarm span shorter than 1,000 bp in accordance with CNV definitions.

### Setting a dynamic sliding window for CNV calling

Conventional CNV detection tools typically segment the genome into fixed window bins (e.g., 1 kb or 5 kb) **[24]**, independently compute coverage or *logR* within each window, and compare these values against preset thresholds to call CNVs (Figure 3A and B). However, the fixed-window strategy suffers from two major drawbacks: If the window is too large, signals from small CNVs can be diluted into adjacent normal windows, causing false negatives; if it is too small, boundary regions may generate false positives and reduce resolution. Both issues ultimately blur CNV boundary definitions and lead to incorrect or missed calls.

**Figure 3.**
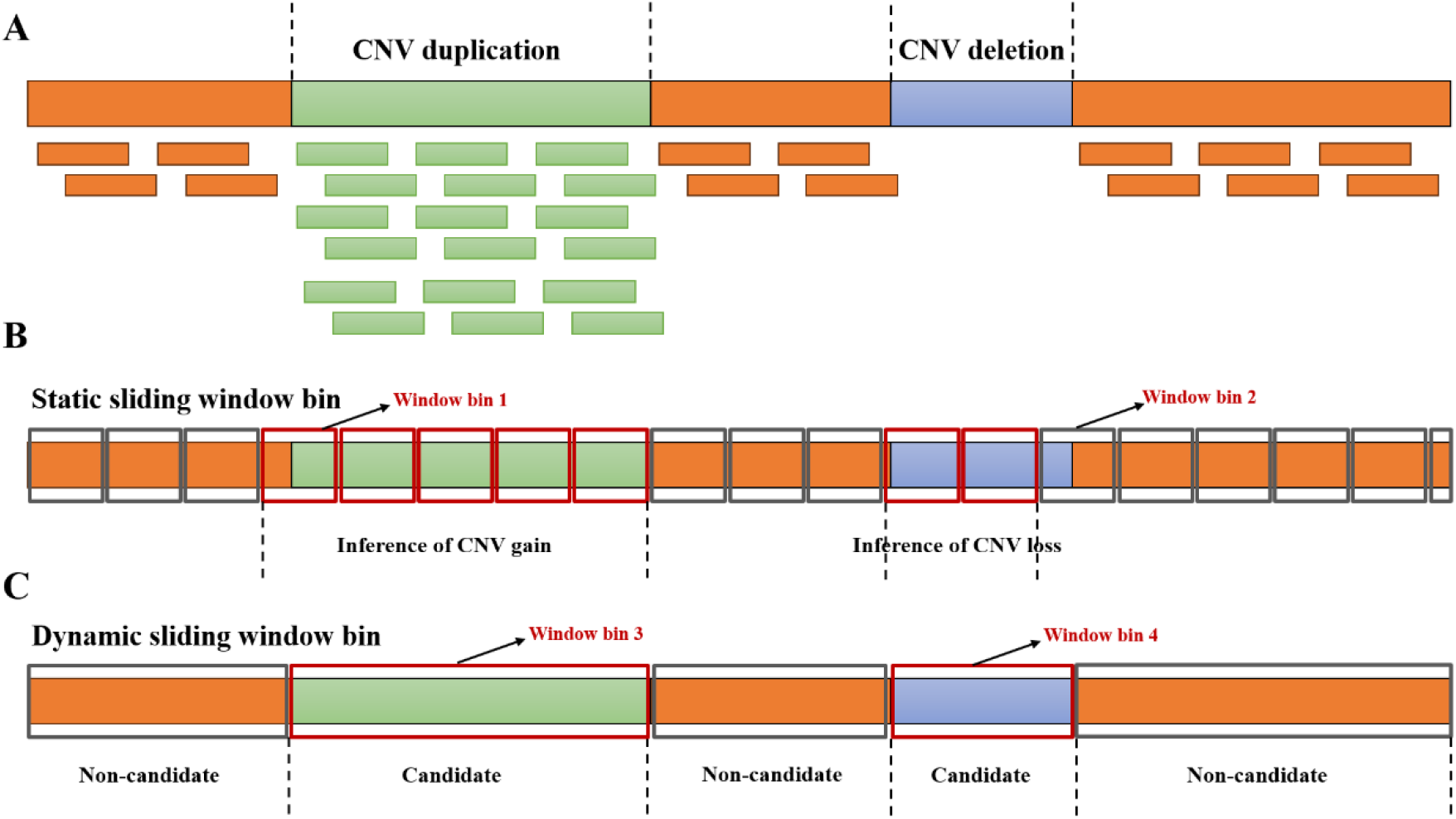
Comparison between the fixed window strategy and dynamic sliding-window strategy. A. Light green and light blue represent copy number gain and loss regions, respectively, in sWGS data. B. Traditional CNV detection tools typically divide the genome into fixed sliding windows and independently calculate coverage or *logR* within each window. CNV signals in Window 1 can cause false positives in adjacent normal windows, while the CNV signal in Window 2 may be diluted by neighboring normal regions, resulting in false negatives. C. Sliding-window strategy of ZIPcnv defines candidate and non-candidate CNV regions based on alarm signals, enabling precise detection of CNVs of varying sizes.

We propose using a dynamic sliding window approach that differs from existing methods. Based on candidate and non-candidate CNV regions, we can set window sizes that depend on the size of the region (Figure 3C). We then calculate the *logR* for each region (either candidate or noncandidate CNVs) within the window bin. Let *s* and *e* represent the starting and ending positions of the region, respectively. Let *x*_*i*_ denote the sequencing depth after correcting base-level coverage at the *i*-th position in the sample, and *p*_*i*_ denote the expected sequencing depth correcting baselevel coverage at the *i*-th position in the reference sample. The *logR* value for this window is defined as:

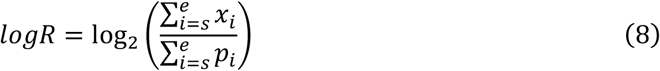

We classify each window according to a fixed threshold of ±0.3:

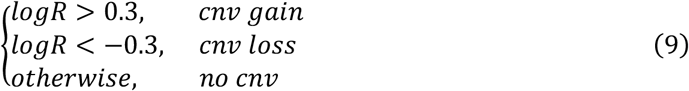

Compared to the traditional fixed window strategy, this dynamic process facilitates subsequent bias correction and region segmentation, and helps accurately detect CNVs of different sizes simultaneously.

## Results

We developed ZIPcnv, a read-depth–based CNV detection method specifically designed for sWGS data that adjusts its window size to accommodate CNVs of varying lengths. We evaluated the performance of ZIPcnv at different coverage levels using both simulated and real-word datasets and compared it against six widely used CNV tools: CNVnator **[25]**, CNVkit **[22]**, PEcnv **[16]**, cn.MOPS **[8]**, QDNAseq **[9]**, and WisecondorX **[10]**. CNVnator and CNVkit are renowned for their overall performance, whereas QDNAseq, cn.MOPS, and WisecondorX are tailored for sWGS applications. PEcnv, a dynamic sliding-window tool of our own design, was also included. In all comparisons, each tool was run with its default parameters, and we assessed sensitivity, precision, and F1 score. CNV calls were considered matched if they overlapped by at least 50% **[26]**. Detailed descriptions and parameter settings for these tools are provided in Supplement 5.

### Study participants and data production

We evaluated ZIPcnv on both simulated and real-world datasets. For the simulated data, we assessed its accuracy at three different sequencing depths. As shown in Figure 4A, we used GSDcreator **[27]** to generate three cohorts of whole-genome samples at 3×, 1×, and 0.1× coverage. Each cohort comprised 1,000 samples—950 containing a single CNV and 50 controls with no CNVs. To simplify downstream analysis, each CNV sample harbored exactly one variant at a random genomic location, with CNV lengths drawn uniformly between 10 kb and 10 Mb. All simulated reads were 35 bp in length with an insert size of 150 bp, mirroring the parameters of real NIPT data.

**Figure 4.**
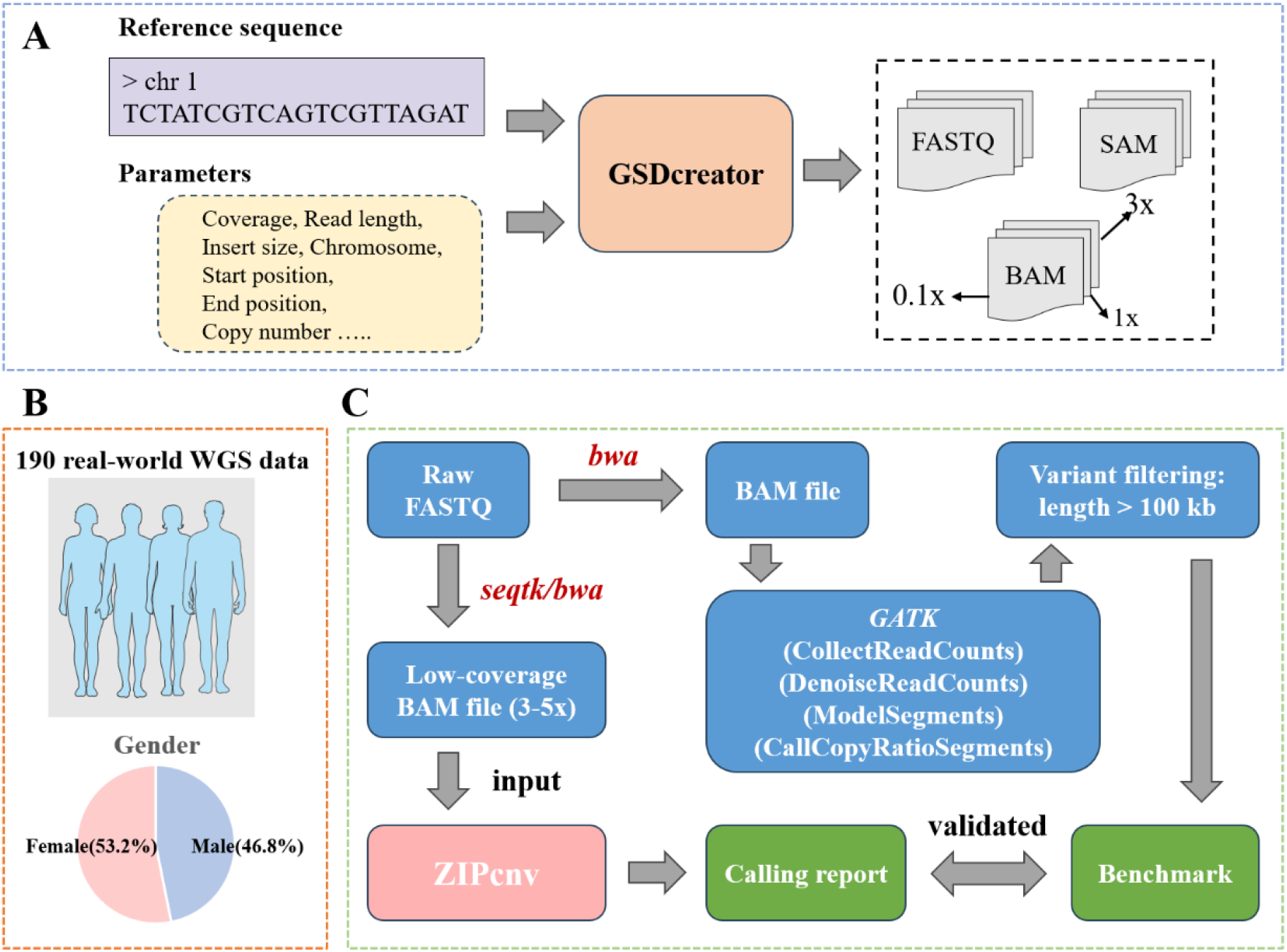
Study participants and data production. A. Workflow for generating simulated data. B. Overview of the BGI-WGS dataset. C. Preprocessing Workflow of BGI-WGS Data

We also collected 190 real whole-genome sequencing samples from Beijing Genomics Institution (BGI) in Shenzhen, China (BGI-WGS). The samples of BGI-WGS were drawn from Han Chinese individuals aged 20–75 years, comprising 89 males (46.8%) and 101 females (53.2%) (Figure 4B). Collected between July 2023 and January 2024, each sample consisted of 5 mL of peripheral blood used for both antibody assays and whole-genome sequencing, with a sequencing depth of approximately 30×–40×. During preprocessing, we employed the GATK CNV discovery pipeline on the raw reads and designated CNVs longer than 100 kb as the benchmark (Figure 4C). The BGI-WGS contained a total of 48,132 CNVs—21,172 gains and 26,960 losses—of which 31,839 ranged from 100 kb to 1 Mb and 16,293 exceeded 1 Mb (Table 2). Finally, we down-sampled each sample to a random target coverage of 3×–5× using *seqtk* and aligned the reads to the GRCh38 reference genome.

**Table 2.**
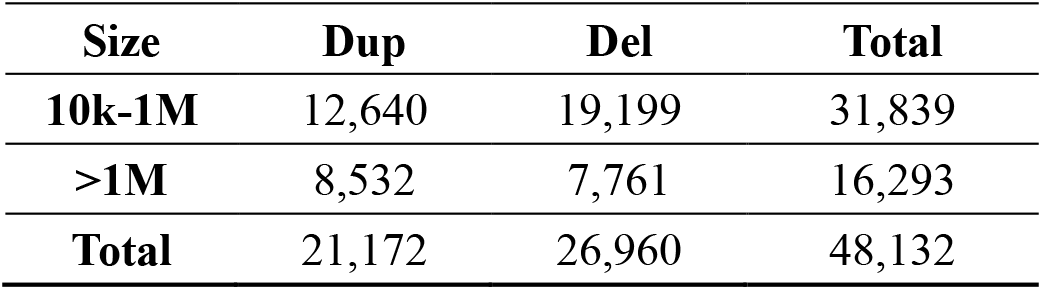
Overview of the number of CNVs in the BGI-WGS.

| Size | Dup | Del | Total |
| --- | --- | --- | --- |
| 10k-1M | 12,640 | 19,199 | 31,839 |
| >1M | 8,532 | 7,761 | 16,293 |
| Total | 21,172 | 26,960 | 48,132 |

### Performance of ZIPcnv at different coverage levels

We evaluated the performance of ZIPcnv on simulated sequencing datasets at varying coverage depths and compared it with several widely used CNV detection methods. Figure 5A-C shows the sensitivity, precision, and F1 score for each tool on the simulated dataset at various low-coverage levels (Table 3). Across all CNV tools, sensitivity, precision, and F1 score declined as coverage decreased, highlighting the increased difficulty of CNV detection at low depths (Table S1-S3, Figure 5D). Traditional CNV tools, including *CNVnator, CNVkit*, and *PEcnv*, exhibit high false-positive rates at 0.1× coverage, with average sensitivity of 0.608 and average precision of 0.447. In contrast, CNV tools tailored for sWGS data—cn.MOPS, QDNAseq, and WisecondorX—delivered overall performance markedly superior to that of traditional CNV tools. At 3× coverage, ZIPcnv delivered on-par performance with other tools, but its advantages became increasingly clear as coverage declined—most notably at 0.1×, where it maintained high sensitivity (0.903) and precision (0.744), underscoring its robustness and reliability.

**Table 3.** Performance comparison between ZIPcnv and other CNV tools on simulated datasets at various low-coverage levels.

| Coverage | 3X |  |  | 1X |  |  | 0.1X |  |  |
| --- | --- | --- | --- | --- | --- | --- | --- | --- | --- |
| Tools | <sup>1</sup>Sens | <sup>2</sup>Prec | <sup>3</sup>F1 | Sens | Prec | F1 | Sens | Prec | F1 |
| *CNVnator* | 0.973 | 0.853 | 0.909 | 0.891 | 0.674 | 0.767 | 0.783 | 0.538 | 0.638 |
| *CNVkit* | 0.879 | 0.779 | 0.826 | 0.665 | 0.557 | 0.606 | 0.513 | 0.405 | 0.453 |
| *PEcnv* | 0.867 | 0.750 | 0.804 | 0.678 | 0.522 | 0.590 | 0.527 | 0.398 | 0.454 |
| *cnMOPS* | 0.855 | 0.868 | 0.862 | 0.709 | 0.697 | 0.703 | 0.422 | 0.369 | 0.394 |
| *QDNaseq* | 0.952 | 0.865 | 0.906 | 0.936 | 0.810 | 0.868 | 0.873 | 0.569 | 0.689 |
| *WisecondorX* | 0.959 | 0.882 | 0.919 | 0.921 | 0.749 | 0.826 | 0.889 | 0.678 | 0.770 |
| *ZIPcnv* | 0.983 | 0.936 | 0.959 | 0.961 | 0.855 | 0.905 | 0.903 | 0.744 | 0.816 |

<sup>1</sup>Sens, Sensitivity; <sup>2</sup>Prec, Precision; <sup>3</sup>F1, F1 score.

**Figure 5.**
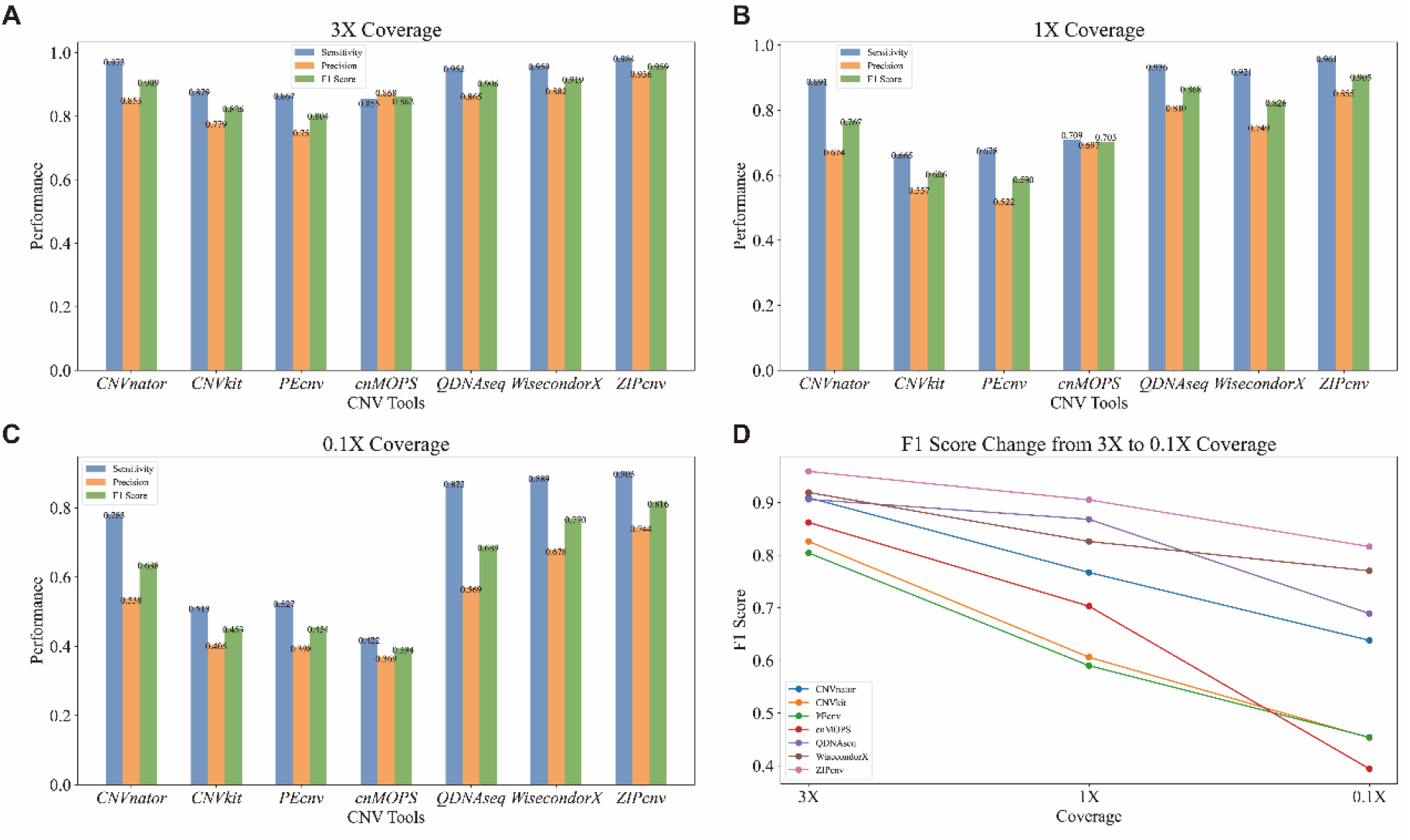
Performance of ZIPcnv and other CNV detection tools across different sequencing coverages. A. Performance at 3X coverage. B. Performance at 1X coverage. C. Performance at 0.1X coverage. D. F1 score change from 3X to 1X coverage.

It should be noted that CNV tools designed specifically for sWGS data often sacrifice some precision to achieve higher sensitivity for small CNVs, albeit with a modest increase in falsepositive rates **[26]**. By employing a dynamic sliding-window strategy, ZIPcnv outperformed alternative tools across all coverage levels in sWGS data, combining high sensitivity with effective reduction of false positives and demonstrating significant potential for real-world applications.

### Performance of ZIPcnv on BGI-WGS

We further evaluated ZIPcnv on BGI-WGS dataset and compared its performance with other CNV detection tools. On BGI-WGS, ZIPcnv achieved an overall sensitivity of 0.935, precision of 0.778, and F1 score of 0.849, outperforming the competing tools (Table 4). Although the precision of ZIPcnv (0.778) was slightly lower than that of cn.MOPS (0.804), its sensitivity substantially exceeded cn.MOPS (Figure 6A). CNVkit reported the highest false-positive count (20,886 events) (Table S4) and a precision of only 0.644. Due to differences in coverage distribution, our previously developed tool PEcnv also underperformed, reaching sensitivity and precision values of just 0.759 and 0.648, respectively.

**Table 4.** Performance comparison between ZIPcnv and other CNV tools on BGI-WGS.

| Tools | Sensitivity | Precision | F1 score |
| --- | --- | --- | --- |
| *CNVnator* | 0.843 | 0.714 | 0.773 |
| *CNVkit* | 0.787 | 0.644 | 0.708 |
| *PEcnv* | 0.759 | 0.648 | 0.699 |
| *cnMOPS* | 0.800 | 0.804 | 0.802 |
| *QDNaseq* | 0.884 | 0.740 | 0.806 |
| *WisecondorX* | 0.919 | 0.753 | 0.828 |
| *ZIPcnv* | 0.935 | 0.778 | 0.849 |

**Figure 6.**
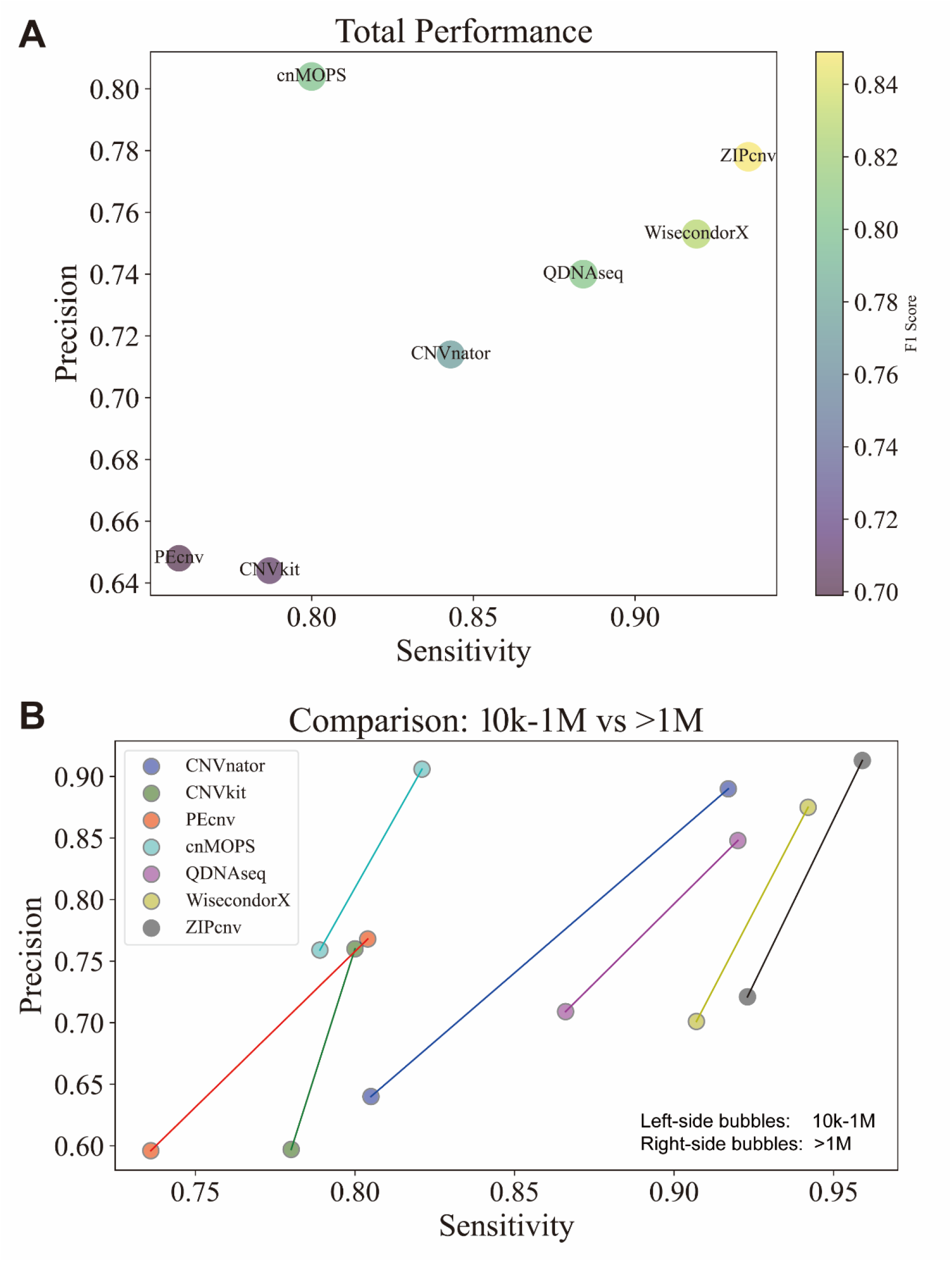
Performance of ZIPcnv and Other CNV Detection Tools on BGI-WGS. A. Overall performance comparison of seven CNV detection tools. The x-axis represents sensitivity, the y-axis represents precision, and the size and color of each bubble indicate the F1 score. B. Performance of the seven CNV detection tools stratified by CNV size: 10 kb–1 Mb and >1 Mb.

For CNVs larger than 1 Mb, ZIPcnv recorded a sensitivity of 0.959, precision of 0.913, and F1 score of 0.935, with only 674 false negatives—leading all tools in this size category (Table S4). However, the gap was narrow: WisecondorX ranked second with sensitivity of 0.942, precision of 0.875, and F1 score of 0.907. CNVnator also demonstrated high sensitivity (0.917) and precision (0.890) for CNVs > 1 Mb (Table 5). Nonetheless, most existing CNV tools find it challenging to detect small CNVs effectively in sWGS data, and their precision and F1 scores generally decrease as CNV size diminishes

**Table 5.** The performance of CNV tools on different size CNVs in the BGI-WGS.

| Size | 10k-1M |  |  | >1M |  |  |
| --- | --- | --- | --- | --- | --- | --- |
| Tools | Sensitivity | Precision | F1 score | Sensitivity | Precision | F1 score |
| *CNVnator* | 0.805 | 0.640 | 0.713 | 0.917 | 0.890 | 0.904 |
| *CNVkit* | 0.780 | 0.597 | 0.676 | 0.800 | 0.760 | 0.780 |
| *PEcnv* | 0.736 | 0.596 | 0.659 | 0.804 | 0.768 | 0.786 |
| *cnMOPS* | 0.789 | 0.759 | 0.773 | 0.821 | 0.906 | 0.862 |
| *QDNaseq* | 0.866 | 0.692 | 0.769 | 0.920 | 0.848 | 0.883 |
| *WisecondorX* | 0.907 | 0.701 | 0.791 | 0.942 | 0.875 | 0.907 |
| *ZIPcnv* | 0.923 | 0.721 | 0.810 | 0.959 | 0.913 | 0.935 |

In the 10 kb–1 Mb range, ZIPcnv achieved sensitivity of 0.923, precision of 0.721, and F1 score of 0.810, with 11,371 false positives and 2,473 false negatives (Table S4). Although its sensitivity and precision decreased by 0.036 and 0.192 compared to the > 1 Mb category, ZIPcnv still demonstrated robust detection capability. In contrast, the precision of CNVnator plummeted to 0.640—a drop of 0.250—while its false positives increased from 1,838 (> 1 Mb) to 14,449, markedly reducing its reliability. Meanwhile, the sensitivity and precision of the other tools also declined to varying degrees (Figure 6B).

Overall, the exceptional performance of ZIPcnv on the BGI-WGS underscores its potential for practical applications, especially in genomics studies that demand high sensitivity and stability.

## Discussion and conclusion

Precise identification of small or low-frequency CNVs in sWGS data remains a major challenge, primarily due to weak signals and high background noise. As previously reported **[28]**, these factors substantially impede the sensitivity and accuracy of CNV detection. In this study, we introduce ZIPcnv, a novel CNV detection tool tailored for sWGS data. ZIPcnv represents a CUSUM control chart to genomic CNV detection, and is particularly effective for low-coverage sequencing data. Compared to the existing tools, ZIPcnv employs statistically dynamic windows and a refined CUSUM accumulation strategy to significantly improve both sensitivity and precision in CNV detection.

ZIPcnv addresses the fundamental limitations of sWGS-based CNV detection by robustly distinguishing true CNV signals from pervasive zero-inflated noise, even under ultra-low coverage conditions. The dynamic adaptation of window sizes enables the detection of CNVs ranging from small focal alterations to large chromosomal segments, thus meeting the increasing demand for comprehensive structural variation analysis in biomedical research. Notably, the superior performance of ZIPcnv in identifying large-segment CNVs further supports its utility for applications requiring high-resolution structural variant profiling, such as disease diagnosis, genetic screening, and personalized treatment planning.

Although ZIPcnv demonstrates excellent performance, several areas warrant further refinement. First, exclusive dependence on RD signals constrains the capacity of ZIPcnv to resolve complex structural rearrangements—such as inversions and translocations—that cannot be accurately distinguished by RD-based approach alone **[29,30]**. Second, ZIPcnv depends heavily on the accuracy and representativeness of the reference baseline. Suboptimal baseline selection may compromise detection precision. Future work should therefore focus on optimizing data-normalization protocols to enhance robustness and interoperability across diverse experimental conditions and sequencing platforms. In addition, extensive validation using clinical cohorts from different disease contexts will be essential to fully establish the utility of ZIPcnv in clinical diagnostics.

In summary, ZIPcnv provides a new analytical framework for interpreting genome data characterized by substantial biological noise and complexity.

## Supporting information

Table S2: Performance comparison between ZIPcnv and other CNV tools on simulated data at coverage 1x.

Table S3:Performance comparison between ZIPcnv aTable S2: Performance comparison between ZIPcnv and other CNV tools on simulated data at coverage 1x.

Table S4: Performance comparison between ZIPcnv

Table S1:Performance comparison between ZIPcnv and other CNV tools on simulated data at coverage 3x.

## Data Availability Statement

The usage of ZIPcnv, related code, scripts, and tools of ZIPcnv are available at https://github.com/Nevermore233/ZIPcnv.

## Author contributions

J.W., X.J. and Z.X. conceived and designed this research; J.W. and Z.X. designed the model; Z.X., J.Y., J.Z. and Y.W. implemented the program and performed the experiments; J.Z., J.Y. and Z.X. provided and analyzed the data. Z.X., J.W. wrote the manuscript. Z.X., J.W., T.W., X.L., L.W., and H.Z. conducted the revision. All authors have read and agreed to the latest version of the manuscript.

