## Supplement 1-5 for "ZIPcnv: accurate and efficient inference of copy number variations from shallow whole-genome sequencing"

**Supplementary Materials**

**Supplement 1. Read depth normalization**

Due to the potential differences in sequencing platforms or sequencing coverage among shallow whole-genome sequencing (sWGS) data, it is necessary to normalize or standardize the initial read depth signals to eliminate inter-platform discrepancies. In this study, we use the median coverage depth of sequencing loci in each sample to remove systematic differences caused by varying sequencing library sizes, and introduce the cross-sample mean coverage depth for each region to correct for sequencing biases inherent to different regions (such as GC content, mappability, repetitive sequences). The normalization formula is as follows:

$$\begin{aligned} \frac{R_{ij}}{R_{j}^{median}}=\frac{R_{ij}^{*}}{R_{i,\cdot}^{mean}}, j=1,2,\ldots,s;i=1,2,\ldots,m\#\left( 1 \right) \end{aligned}$$

where *s* represents the number of sequencing samples, *m* represents the total number of genomic segments in each sample, $R_{ij}$ denotes the read depth signal of the $i$-th segment in the $j$-th sample, $R_{j}^{median}$ is the median read depth signal across all segments in the $j$-th sample, $R_{i,\cdot}^{mean}$ is the mean read depth signal of the $i$-th segment across all samples, and $R_{ij}^{*}$ is the normalized read depth signal. The normalized read depth signal matrix ${{[R}_{ij}^{*}]}_{m*s}$ can be obtained using this formula. The final output are standardized files (.json) corresponding to each sample.

**Supplement 2. Reference baseline establishment**

We established a reference baseline for the CNV calling. CNV detection tools based on high-coverage sequencing typically require only a single normal sample as a control to provide a sufficient signal-to-noise ratio. However, in low-coverage sequencing, the higher level of random noise makes it difficult for a single control sample to effectively distinguish between noise and true copy number variations, which could compromise detection accuracy. Therefore, following Raman's approach [1], we selected at least 50 normal samples as the baseline, with the read depth at each locus being the average of these 50 samples. The final output is a reference baseline file for each chromosome (.npz).

**Supplement 3. From ZIP distribution to normal distribution: transformation and theoretical proof.**

Original sWGS data typically exhibit zero-inflation characteristics, meaning that a substantial number of genomic regions have zero read depth. The Zero-Inflated Poisson (ZIP) distribution effectively models this phenomenon, capturing a large proportion of data points as structural zeros, while the remaining non-zero observations follow a Poisson distribution [2]. Specifically:

$$\begin{aligned} P\left( X=x \right)=\left\{ \begin{matrix} \pi+\left( 1-\pi\right)e^{-\lambda}, & x=0 \\ \left( 1-\pi\right)\frac{e^{-\lambda}\lambda^{x}}{x!}, & x>0 \end{matrix} \right.\#\left( 2 \right) \end{aligned}$$

Here, $\pi$ represents the proportion of structural zeros, and $\lambda$ is the rate parameter of the Poisson process. The expectation and variance of the ZIP distribution can be expressed as:

$$\begin{aligned} E\left( X \right)=\left( 1-\pi\right)\lambda\#\left( 3 \right) \end{aligned}$$

$$\begin{aligned} Var\left( X \right)=\left( 1-\pi\right)\lambda\left( 1+\pi\lambda\right)\#\left( 4 \right) \end{aligned}$$

Through sliding window averaging, the data is smoothed, and the original zero-inflated Poisson characteristics are transformed into a more continuous and symmetric distribution. According to the Central Limit Theorem (CLT), as the window size increases, this new distribution will converge to a normal distribution [3]. The reasoning process is as follows.

Consider a fixed window containing $n$ genomic loci, where their original sequencing read depths $X_{i}$ are denoted as random variables:

$$\begin{aligned} X_{1}, X_{2}, \ldots,X_{n}\#\left( 5 \right) \end{aligned}$$

Assume that these random variables are independent and identically distributed (or at least approximately independent and identically distributed), and each random variable follows a ZIP distribution. We are interested in the average of these random variables:

$$\begin{aligned} \bar{X}=\frac{1}{n}\sum_{i=1}^{n} X_{i}\#\left( 6 \right) \end{aligned}$$

The expectation of the $\bar{X}$ is:

$$\begin{aligned} E\left( \bar{X} \right)=E\left( \frac{1}{n}\sum_{i=1}^{n} X_{i} \right)=\frac{1}{n}\sum_{i=1}^{n} E\left( X_{i} \right)=\frac{1}{n}\times n\cdot\left( 1-\pi\right)\lambda=\left( 1-\pi\right)\lambda\#\left( 7 \right) \end{aligned}$$

The variance of the $\bar{X}$ is:

$$\begin{aligned} Var\left( \bar{X} \right)=Var\left( \frac{1}{n}\sum_{i=1}^{n} X_{i} \right)=\frac{1}{n^{2}}\sum_{i=1}^{n} Var\left( X_{i} \right)=\frac{1}{n^{2}}\cdot n\cdot\left( 1-\pi\right)\lambda\left( 1+\pi\lambda\right)\#\left( 8 \right) \end{aligned}$$

So:

$$\begin{aligned} Var\left( \bar{X} \right)=\frac{\left( 1-\pi\right)\lambda\left( 1+\pi\lambda\right)}{n}\#\left( 9 \right) \end{aligned}$$

According to the CLT, when $n$ is large, regardless of the original distribution of $X_{i}$, as long as the expectation and variance are finite, the following holds:

$$\begin{aligned} \bar{X}\underset{\to}{d}N\left( \left( 1-\pi\right)\lambda, \frac{\left( 1-\pi\right)\lambda\left( 1+\pi\lambda\right)}{n} \right)\#\left( 10 \right) \end{aligned}$$

Therefore, based on the above derivation, the averaged read depth after applying a sufficiently large sliding window follows (or approximately follows) a normal distribution, with a mean of $\left( 1-\pi\right)\lambda$ and a variance of $\frac{\left( 1-\pi\right)\lambda\left( 1+\pi\lambda\right)}{n}$. This not only aligns with the common assumptions in classical CNV detection methods but also provides a solid theoretical foundation for subsequent CUSUM-based statistical procedures that rely on the assumption of normality.

**Supplement 4. Theoretical basis of CUSUM application to sWGS Data**
(1) Basic setup of CUSUM control chart
The CUSUM (Cumulative Sum) control chart is a statistical method used to monitor whether the mean of a data sequence undergoes a continuous small shift [4]. We assume that the observed data sequence is:

$$\begin{aligned} X_{1}, X_{2}, \ldots,X_{n}\#\left( 11 \right) \end{aligned}$$

Under normal conditions (In-Control), these data satisfy:

$$\begin{aligned} X_{i}\sim N\left( \mu_{0}, \sigma^{2} \right)\#\left( 12 \right) \end{aligned}$$

In sWGS data, it is:

$$\begin{aligned} X_{i}\sim N\left( \left( 1-\pi\right)\lambda, \frac{\left( 1-\pi\right)\lambda\left( 1+\pi\lambda\right)}{n} \right)\#\left( 13 \right) \end{aligned}$$

When an abnormality occurs (out-of-control), the distribution of data becomes:

$$\begin{aligned} X_{i}\sim N\left( \mu_{1}, \sigma^{2} \right),\mu_{1}=\mu_{0}+\delta,\delta\neq0\#\left( 14 \right) \end{aligned}$$

Here, $\delta$ represents the shift, which is typically small but persistent.

(2) Statistical construction of the CUSUM control chart
The core idea of the CUSUM control chart is to continuously monitor the cumulative effect of mean shifts by summing the deviations of the observed values from the target mean $\mu_{0}$. Specifically, the CUSUM statistic is defined as:

$$\begin{aligned} C_{t}= \max\left[ 0,C_{t-1} +\left( X_{t}-\mu_{0}-k \right) \right]\#\left( 15 \right) \end{aligned}$$

where $C_{0}=0$ is the initial condition, $k$ is the reference value, typically set to half of the shift, i.e., $k=\delta/2$. The $X_{t}$ is the observed data point. The purpose of introducing the reference value $k$ is to accumulate only when the data shift exceeds a certain threshold, thus making the CUSUM more sensitive to subtle shifts and suppressing the interference of random noise.

(3) Applicable data conditions and statistical properties of the CUSUM statistic

In 1971, G. Lorden proved that under the assumption of data being independent and normally distributed, CUSUM achieves asymptotic optimality in detecting persistent small mean shifts [5]. In other words, among all statistical methods with the same detection sensitivity level, CUSUM minimizes the expected detection delay. A brief proof is given as follows.

Assuming a true shift $\delta>0$, the earliest time the CUSUM statistic Ct triggers an alarm is:

$$\begin{aligned} T_{C}=inf\left\{ t\geq1:C_{t}\geq h \right\}\#\left( 16 \right) \end{aligned}$$

Where $C_{t}$ represents the CUSUM statistic at time point $t$ and $h$ is the alarm threshold of the control chart. Lorden rigorously proved that under the same false alarm probability $\alpha$, as the shift magnitude $\delta$ approaches zero, the asymptotic optimal bound for the expected detection delay $E_{\delta}\left( T_{C} \right)$ of the CUSUM is:

$$\begin{aligned} \lim_{\delta\to0} \frac{E_{\delta}\left( T_{C} \right)}{\left| \log\left( \alpha\right) \right|}=\frac{2\sigma^{2}}{\delta^{2}}\#\left( 17 \right) \end{aligned}$$

Here, $\frac{2\sigma^{2}}{\delta^{2}}$ represents the asymptotic optimal bound for the expected detection delay $E_{\delta}\left( T_{C} \right)$. As the shift magnitude $\delta$ approaches zero, any detection method must significantly increase its expected detection delay $E_{\delta}\left( T_{C} \right)$ to maintain a fixed low false alarm probability $\alpha$. However, among all control charts, when the shift is small and persistent, the expected detection delay $E_{\delta}\left( T_{C} \right)$ for CUSUM is proportional to $\frac{1}{\delta^{2}}$, making it the slowest among all control charts [5].

In sWGS data, the original read depth follows a zero-inflated Poisson distribution. By applying a large sliding window strategy, the averaged sWGS data satisfy the two core conditions required for the CUSUM statistic: (I) the data are approximately normally distributed, and (II) anomalies (CNVs) manifest as persistent and stable small shifts. Therefore, we adapt the CUSUM control chart, originally developed for quality control, to CNV detection. This not only demonstrates the transferability of interdisciplinary tools but also highlights the inherent "process fluctuation" nature of copy number variation detection.

**Supplement 5. The descriptions and parameter settings of CNV detection tools**

We evaluated the performance of ZIPcnv using both simulated and real-world data under different coverage levels and varying CNV lengths, and compared it with several widely used CNV detection tools, including *CNVnator* [6], *CNVkit* [7], *PEcnv* [8], *cn.MOPS* [9], *QDNAseq* [10], and *WisecondorX* [1].

*CNVnator* is a CNV detection tool based on read depth. It creates fixed-size windows across the genome, calculates the read depth for each window, and identifies CNV events based on changes in depth. Due to its fast computation speed and low memory usage, *CNVnator* is widely applied in CNV analysis of whole-genome sequencing data. In our experiments, the default window size of 5000 was used.

*CNVkit* is a CNV detection tool specifically designed for whole-genome or targeted sequencing data. It infers the copy number state of the genome by extracting coverage signals from the sequencing data, and is particularly effective in handling cancer data and targeted sequencing applications. In our experiments, the window size was set to 267, with threshold values set to -1.1, -0.4, 0.3 and 0.7.

*PEcnv* is a CNV detection tool based on a dynamic sliding window, primarily used for whole-genome sequencing data. By dynamically adjusting the window size, *PEcnv* can adapt to the varying lengths of variation features in the genome. It performs especially well in handling complex genomic regions, making it suitable for fine-grained CNV analysis. Since it uses a dynamic sliding window, no window size needs to be specified.

*cn.MOPS* is a CNV detection tool based on statistical models, particularly suitable for multi-sample datasets. By detecting shared copy number variations in a set of samples, it significantly reduces the false positive rate caused by technical noise. cn.MOPS is implemented through the R package "cn.mops" and does not require reference samples or parameter adjustments.

*QDNAseq* is designed for shallow whole-genome sequencing (sWGS) data. It reduces the noise caused by low sequencing depth by correcting coverage fluctuations, making it suitable for cancer research and non-invasive prenatal testing (NIPT) scenarios. In our experiments, the R package for *QDNAseq* was installed via the Bioconductor tool, with the window size set to 15,000.

*WisecondorX* is specifically designed for non-invasive prenatal testing (NIPT) and is mainly used for detecting fetal chromosomal abnormalities, including CNVs and aneuploidy, in shallow whole-genome sequencing (sWGS) data. *WisecondorX* requires at least 50 healthy samples as a reference set, with representative male and female samples in a 1:1 ratio for optimal detection accuracy. The workflow consists of three steps: first, converting the aligned reads (bam/cram format) into .npz files; second, creating a reference set; and third, predicting CNVs. It is particularly adept at detecting variations greater than 5MB, with the bin size set to 10k and Z-score cutoff set to 5.
